# Confounds of using the *unc-58* selection marker highlights the importance of genotyping co-CRISPR genes

**DOI:** 10.1101/2021.05.26.445785

**Authors:** Helena Rawsthorne-Manning, Fernando Calahorro, Patricia Gonzalez Izquierdo, Lindy Holden-Dye, Vincent O’Connor, James Dillon

## Abstract

Multiple advances have been made to increase the efficiency of CRISPR/Cas9 editing using the model genetic organism *Caenorhabditis elegans* (*C. elegans*). Here we report on the use of co-CRISPR ‘marker’ genes: worms in which co-CRISPR events have occurred have overt, visible phenotypes which facilitates the selection of worms that harbour CRISPR events in the target gene. Mutation in the co-CRISPR gene is then removed by outcrossing to wild type but this can be challenging if the CRISPR and co-CRISPR gene are hard to segregate. However, outcrossing can be avoided by selecting worms of wild type appearance from a ‘jackpot’ brood. These are broods in which a high proportion of the progeny of a single injected worm display the co-CRISPR phenotype suggesting high CRISPR efficiency. This can deliver worms that harbour the desired mutation in the target gene locus without the co-CRISPR mutation. We have successfully generated a discrete mutation in the *C. elegans nlg-1* gene using this method. However, in the process of sequencing to authenticate editing in the *nlg-1* gene we discovered genomic rearrangements that arise at the co-CRISPR gene *unc-58* that by visual observation were phenotypically silent but nonetheless resulted in a significant reduction in motility scored by thrashing behaviour. This highlights that careful consideration of the hidden consequences of co-CRISPR mediated genetic changes should be taken before downstream analysis of gene function. Given this, we suggest sequencing of co-CRISPR genes following CRISPR procedures that utilise phenotypic selection as part of the pipeline.

## Introduction

Originally identified as a mechanism of bacterial immunity (Mojica *et al*., 2005), CRISPR/Cas9 is now a widely used genome editing technique that allows for precise and customisable DNA modification (Jinek *et al*., 2012). Cas9 endonuclease cleavage of DNA results in a double strand break which can then be repaired via homology directed repair (HDR) or non-homologous end joining (NHEJ) pathways. HDR can be utilised to generate precise DNA edits because it allows for the integration of an exogenous repair template into the gene of interest (Adli, 2018). In contrast, NHEJ can create noncontrolled insertions and deletions into a sequence and is therefore often used to generate disruptive genetic mutations that aim to knockout gene function (Hsu, Lander and Zhang, 2014). The ability to precisely manipulate DNA in this way has seen CRISPR/Cas9 facilitate insight into gene function in multiple model organisms (Ma and Liu, 2015).

*C. elegans* is an attractive model organism for the application of CRISPR/Cas9 because their genetic tractability and short life cycle allows for the rapid generation of progeny with cloned genetic identity (Kaletta and Hengartner, 2006). Systems level analysis of gene function in *C. elegans* means CRISPR mediated modifications can be investigated at a molecular and behavioural level (Rawsthorne *et al*., 2020; McDiarmid *et al*., 2019; Wong *et al*., 2019; Greene *et al*., 2016; López-Cruz *et al*., 2019). Furthermore, due to the genetic homology of mammalian and *C. elegans* genomes (Sonnhammer and Durbin, 1997), CRISPR/Cas9 has been used to investigate specific human disease related mutations (Rawsthorne *et al*., 2020; Wong *et al*., 2019; McDiarmid *et al*., 2019).

Despite its advantages, the efficiency of CRISPR/Cas9 can be limited by a number of factors. For example, the interaction of the single guide RNA (sgRNA) Cas9 complex with target DNA requires Cas9 to bind to a protospacer adjacent motif (PAM) (Dickinson and Goldstein, 2016). It has been shown that the closer the PAM to the target site the better the editing efficiency (Paix *et al*., 2014). However, the prevalence of PAM sites has been shown to be scarce within some *C. elegans* genes (El Mouridi *et al*., 2017) thus limiting the regions of the genome that can be targeted efficiently. A method developed to try and negate this issue involves the transplantation of a d10 sequence into the gene of interest prior to generating the desired edit. The d10 sequence contains a protospacer and PAM from the endogenous *dpy-10 C. elegans* gene. Transplantation of the necessary motifs for efficient CRISPR/Cas9 therefore renders the gene of interest more amenable to editing and hence enhances the scope of CRISPR in *C. elegans* (El Mouridi *et al*., 2017).

Another technique used to increase the efficiency of the CRISPR process in *C. elegans* is the use of co-CRISPR genes (Arribere *et al*., 2014; Kim *et al*., 2014). A co-CRISPR gene is a gene which is subjected to gene editing simultaneously with the target gene of interest. Upon mutation, co-CRISPR target genes result in a visibly obvious, marker phenotype which can be easily distinguished from worms with a wild-type appearance. The marker phenotype provides a visual representation of CRISPR efficiency and therefore minimises the number of progeny that need to be sequenced to identify the desired edit. Multiple co-CRISPR genes have been identified, including *dpy-10, rol-6, sqt-1* and *unc-58* (Arribere et al., 2014; Kim et al., 2014). Mutations in *dpy-10* result in dumpy, roller or dumpy roller phenotypes, depending on the mutant genotype. Mutations in *rol-6* and *sqt-1* result in roller phenotypes causing the worms to move in a circular pattern (Arribere *et al*., 2014). *unc-58* encodes a two-pore domain potassium channel (K2P) in *C. elegans* (Salkoff, 2005) which plays an important role in maintaining neuronal and muscle cell excitability (Salkoff *et al*., 2001; Reiner, Weinshenker and Thomas, 1995). A prime function of the protein encoded by *unc-58* is to act as a leak channel that is important in setting the resting membrane potential (Kasap *et al*., 2018). A gain-of-function (GOF) mutation in *unc-58(e665)* fourth transmembrane helix results in a hypercontracted and uncoordinated phenotype which leaves the worm immobile (Brenner, 1974; Park and Horvitz, 1986). Interestingly, it is hypothesised that this hypercontraction results from a change in the selectivity of the channel from K^+^ to Na^+^ (Kasap *et al*., 2018). This pronounced behavioural change provides a simple binary readout of the integrity of the *unc-58* locus. In particular, the switch from immobile to motile provides an obvious measure of reversion from GOF to WT. The easily recognisable phenotype of *unc-58* GOF mutants has led to its successful widespread use in technologies that are aided by phenotypic markers (Kasap and Dwyer, 2020; Huumonen *et al*., 2012; Hartman *et al*., 2014) and this has recently been extended to co-CRISPR techniques (Arribere *et al*., 2014; El Mouridi *et al*., 2017; Wang *et al*., 2018).

Easily recognisable co-CRISPR phenotypes can also be used to identify so called jackpot broods. Jackpot broods are a population of progeny derived from a single parent, injected with Cas9, sgRNAs and repair templates to facilitate CRISPR at both a target gene and co-CRISPR gene, in which a high percentage of the population displays the co-CRISPR marker phenotype (Paix *et al*., 2015). A large number of co-CRISPR marked progeny provides a visual representation of high efficiency CRISPR editing at the co-CRISPR gene and is therefore indicative of high CRISPR efficiency at the target gene of interest (Arribere et al., 2014; Kim et al., 2014). co-CRISPR marked worms from jackpot broods are therefore selected for sequencing for the mutation of interest. Within jackpot broods there are also siblings that do not display the easily recognisable co-CRISPR phenotype. Whilst it can be assumed, based on the phenotype, that these unmarked siblings are WT for the co-CRISPR loci they can still carry the desired CRISPR mutation in the gene of interest, albeit at a lower frequency than in the progeny that show the co-CRISPR phenotype (Paix *et al*., 2014). It can therefore be beneficial to sequence some apparently wild type siblings from high efficiency jackpot broods for the mutation of interest. For example, it has been suggested that if the gene of interest is located on the same chromosome as the co-CRISPR gene then unmarked siblings should be selected for sequencing to avoid the need to backcross the co-CRISPR mutation, which would be complicated for genes that are chromosomally linked (El Mouridi *et al*., 2017). It is noteworthy that when selecting marked and unmarked progeny, the integrity of the co-CRISPR gene is assumed based on the worms’ phenotype and therefore is rarely sequenced as part of the CRISPR pipeline (El Mouridi *et al*., 2017; Wang *et al*., 2018; Schreier *et al*., 2020).

Recently, a method of CRISPR/Cas9 has been developed that combines multiple techniques to increase the efficiency of CRISPR and allows for highly targeted editing in *C. elegans*. The method encompasses a two-step procedure which uses a d10 sequence to facilitate the generation of an edit in a precise location within the gene of interest and uses co-CRISPR marker genes for rapid phenotypic screening (El Mouridi *et al*., 2017). We utilised this method with the aim of mimicking an autism associated missense variant, R451C in neuroligin (Jamain *et al*., 2003), in order to investigate the mutant in a motility based food leaving assay that scores social behaviour (Rawsthorne *et al*., 2020). This first involved transplantation of a d10 sequence into the *C. elegans nlg-1* gene. Simultaneously, the co-CRISPR gene *unc-58* was edited to generate a GOF mutation that results in an uncoordinated phenotypic marker for visual screening of CRISPR efficiency. Following this, in a second round of CRISPR editing, the d10 sequence in *nlg-1* was replaced by a repair template containing the sequence for the desired R451C mutation. Again, simultaneously another co-CRISPR gene, *dpy-10*, was edited to generate *dpy-10(cn64)* GOF mutant for the purpose of phenotypic screening.

In this study, we describe an unexpected CRISPR mediated mutation that occurred within the co-CRISPR gene *unc-58*. The resulting mutants did not display the hypercontraction characteristic of the *unc-58* GOF but rather resulted in a subtle loss-of-function (LOF) not discernible by visual inspection of gross locomotion. We use this to suggest that the phenotypic appearance of *C. elegans* is not a robust way to determine the genotype of co-CRISPR genes. Given this, we suggest routine sequencing of co-CRISPR genes following CRISPR procedures that utilise phenotypic selection before downstream analysis.

## Materials and Methods

### *C. elegans* culturing and strains used

All *C. elegans* strains were maintained using standard conditions (Brenner, 1974). Strains used: Bristol N2 (wild-type), VC228 *nlg-1(ok259) X* (x6 outcrossed), CB665 *unc-58(e665) X*, TN64 *dpy-10(cn64) II*, provided by *Caenorhabditis* Genetics Center (CGC). XA3780 *nlg-1(qa3780) X* (x2 outcrossed), XA3773 *unc-58(qa3788) X; nlg-1(qa3780) X* (x1 outcrossed), XA3788 *unc-58(qa3788) X* (x2 outcrossed), generated in this study. JIP1154 *unc-58(bln223) X*, provided by Thomas Boulin.

### CRISPR/Cas9 genome editing

The previously reported method by El Mouridi et al was followed for CRSIPR/Cas9 editing (El Mouridi *et al*., 2017). L4+1 day old hermaphrodites were microinjected with expression vectors for Cas9 and sgRNAs targeting *nlg-1* and *unc-58* or *dpy-10* for the first and second round of CRISPR, respectively. The design of the sgRNAs and repair templates to edit *unc-58, dpy-10* and *nlg-1* genes were as previously reported (Rawsthorne *et al*., 2020; Arribere *et al*., 2014). sgRNAs and repair templates were purchased from Integrated DNA Technologies (IDT^™^ – Integrated DNA Technologies). All injection reagents were diluted in molecular grade water to a final concentration of 50ng/µl in the CRISPR mix.

### Molecular Screening and sequencing

*nlg-1* primers, forward 5’-ATGAGTATACAGATTGGGAAAATCCC-3’ and reverse 5’-ACTGTTTGGTTGCTCTTGGCTCCAAG-3’, were used to amplify the CRISPR targeted region of *nlg-1* using a single worm PCR protocol (He, 2011). A BanI site was used to screen PCR amplicons from individual worms for the incorporation or loss of the d10 sequence in *nlg-1*. The primers used for sequencing of the target regions of *unc-58* and *dpy-10* were: forward 5’-GACTCGGAGATATCGTTGTGACTG-3’, reverse 5’-CGCGGAGTTCGTTATCCAGGAAG-3’ and forward 5’-ACTAATTCAGAGTCATCATCTCGCC-3’, reverse 5’-CATCAATTCCCTTAAGTCCTGGTGG-3’ respectively.

### Removal of *unc-58* background mutation

*unc-58(qa3788); nlg-1(qa3780)* double CRISPR mutant strain was backcrossed with N2 males once to generate heterozygous F1 progeny. These progeny were cloned by picking a single F1 onto individual plates before being left for 2-3 days to self-fertilise and F2 progeny were screened using a single worm PCR protocol (He, 2011) and restriction digest. Restriction digest of *unc-58 and nlg-1* PCR amplicons were performed using restriction enzymes BsiWI and BstBI respectively to screen for the presence or absence of CRISPR mediated edits. Restriction enzymes were supplied by New England BioLabs (NEB) and used according to manufacturer instructions.

### Food leaving assay

Food leaving assays were carried out as previously described (Rawsthorne *et al*., 2020). Briefly, 50µl of OP50 *E. coli* at OD_600_ of 0.8 was gently spotted on to the middle of an unseeded plate the day prior to the assay. Seven age synchronised L4+1 day old hermaphrodites were gently picked onto the centre of the bacterial lawn on the assay plate. At 2 and 24 hours food leaving events were counted visually using a Nikon SMZ800 microscope (X10 magnification) during 30 minute observations. A food leaving event was defined as when the whole of the worm’s body exited the bacterial lawn. N2 animals were used as a paired control, run in parallel with the strain under investigation and the investigator was blind to the genotypes being observed.

### Thrashing assay

Using a 24 well plate, a single worm was picked per well containing 500µl of M9 with 0.1% bovine serum albumin and left for 5 minutes before thrashing was measured. For each worm, thrashing events were visually counted under a Nikon SMZ800 microscope (X30 magnification) for a period of 30 seconds. This was repeated three consecutive times and the mean was calculated. Each thrash was defined as a complete movement through the midpoint of the worms body and back. N2 animals were used as a paired control, run in parallel with the strain under investigation and the investigator was blind to the genotypes.

### *unc-58* protein sequence analysis

UNC-58, isoform b protein sequence was downloaded from WormBase version WS278. The wild-type or CRISPR mutant protein sequence was entered into the membrane topology prediction tool TMHMM (v.2.0) (Sonnhammer, von Heijne and Krogh, 1998; Krogh *et al*., 2001) to determine the predicted transmembrane topology.

## Results

### The use of co-CRISPR genes facilitated the generation of a precise *nlg-1* edit

We used co-CRISPR genes *unc-58* and *dpy-10* to facilitate the generation of a precise edit in *C. elegans nlg-1* gene. 84 N2 animals were injected with plasmids encoding Cas9 and a sgRNA and repair template designed to transplant a d10 sequence (El Mouridi *et al*., 2017) into the gene of interest, *nlg-1*. In addition, the CRISPR injection mix included a sgRNA and repair template designed to generate a GOF mutation in the co-CRISPR gene *unc-58* (Figure 1A). This was used as a CRISPR efficiency marker based on the hypercontraction and uncoordinated phenotype that results from the mutation. 7 of the 84 parental injected worms produced F1 broods containing progeny that displayed an uncoordinated phenotype, supporting the notion that the *unc-58* gene had been successfully edited (Figure 1B). 2 of these 7 broods were identified as jackpot broods, containing more than 30 progeny with an *unc-58* paralysis phenotype. In total, 72 *unc-58* marked progeny were selected for molecular screening and sequencing of the target region of the *nlg-1* gene (Figure 1A). We also selected 100 unmarked, apparently wild type, siblings from these jackpot broods for screening because *nlg-1* and *unc-58* are both located on the same chromosome (X) and approximately 3.5mb apart. Identifying the *nlg-1* edit in an unmarked worm can be beneficial because it avoids the need for backcrossing to separate two chromosomally linked edits. This resulted in the identification of a single *nlg-1* d10 entry strain in an unmarked F1 worm from a jackpot brood which displayed no obvious *unc-58* marked phenotype. Consistent with other studies, the lack of *unc-58* marked phenotype was used to assume that the *nlg-1* d10 entry strain was WT for the *unc-58* co-CRISPR loci (El Mouridi *et al*., 2017; Wang *et al*., 2018; Schreier *et al*., 2020).

**Figure 1:**
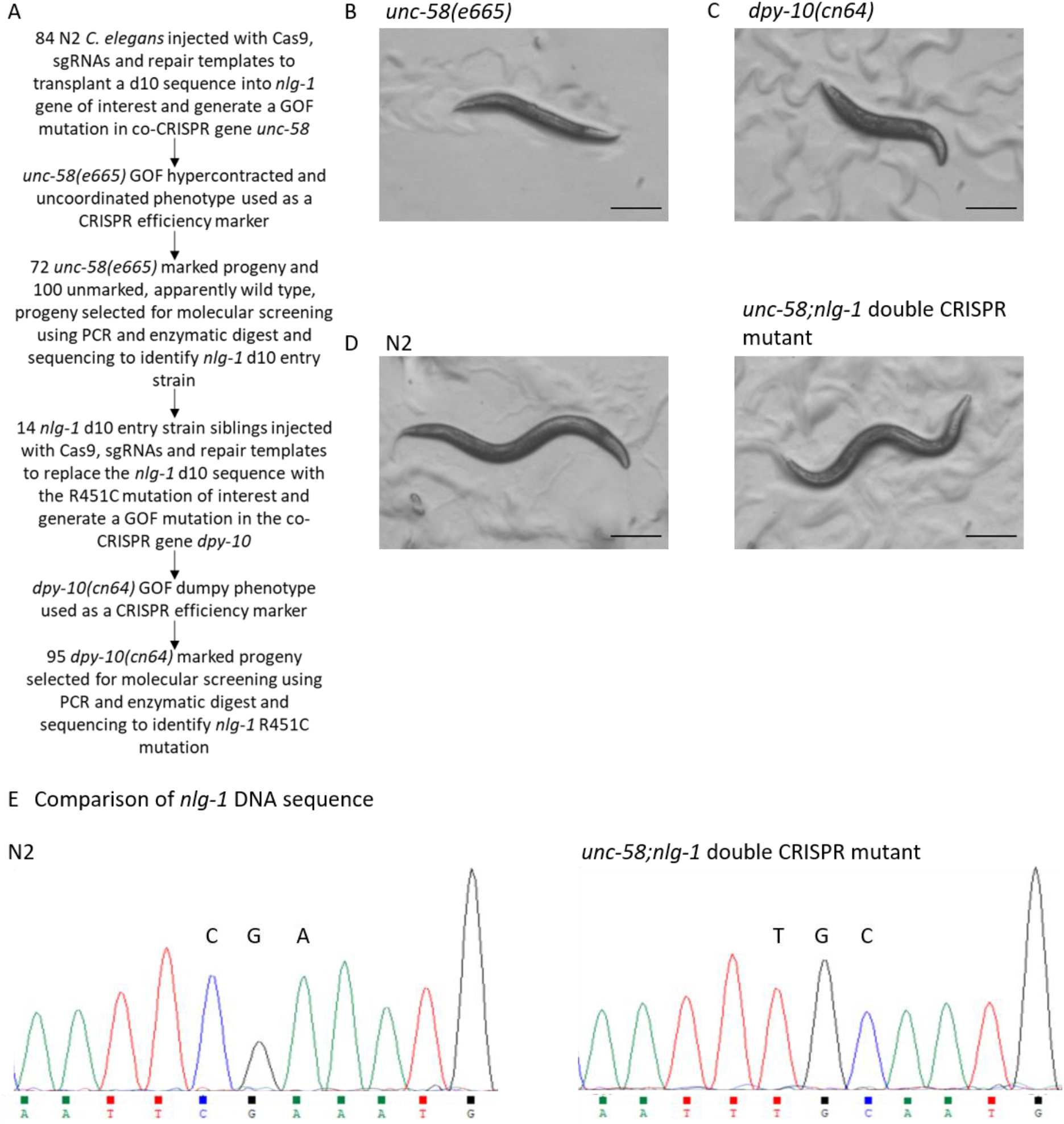
co-CRISPR genes used to facilitate the generation of a *nlg-1* CRISPR mutant. **A** Flow diagram summarising the steps taken to generate a CRISPR/Cas9 mutant which utilised two co-CRISPR genes. **B** Representative image of an *unc-58(e665)* GOF *C. elegans* mutant strain which has an uncoordinated phenotype with severe motility deficits which result in sluggish movement. **C** Representative image of a *dpy-10(cn64)* GOF *C. elegans* mutant strain displaying a dumpy phenotype. **D** Representative images of N2 wild-type and *unc-58(qa3788);nlg-1(qa3780)* double CRISPR mutant strain that highlight the gross phenotypic similarities between the two strains. **B-D** All images show L4+1 day old hermaphrodites and were taken at 30x magnification. All scale bars represent 0.2mm. **E** Chromatograms showing partial *nlg-1* DNA sequences from N2 and *unc-58(qa3788);nlg-1(qa3780)* double CRISPR mutant strain. The codons of interest, CGA and TGC are indicated. TGC encodes the desired *nlg-1* R451C mutation. DNA sequences are shown in the 5’ to 3’ orientation.

Next, 14 d10 entry siblings, from the identified *nlg-1* d10 entry strain, were injected with plasmids encoding Cas9, a sgRNA and a repair template designed to replace the *nlg-1* d10 sequence with the R451C mutation of interest. In addition, the CRISPR injection mix included a sgRNA and repair template designed to generate a GOF mutation in the co-CRISPR gene *dpy-10* (Figure 1A). Four of the injected parental worms produced *dpy-10* marked F1 progeny, displaying a mixture of dumpy (Figure 1C), roller and dumpy roller phenotypes. Due to the high efficiency of the *dpy-10* sgRNA previously reported (El Mouridi *et al*., 2017), the frequency of *dpy-10* marked progeny was higher than that of *unc-58* marked progeny (El Mouridi *et al*., 2017). We identified three plates containing 9-15 *dpy-10* marked progeny each and one plate with 56 marked progeny. Unlike *unc-58, dpy-10* is not located on the same chromosome as *nlg-1* so we selected only *dpy-10* marked progeny for molecular screening. In total, 95 *dpy-10* marked progeny were selected for molecular analysis and sequencing of the target region of the *nlg-1* gene. We identified one dumpy phenotype worm that contained the R451C mutation in *nlg-1*. The aim for the *nlg-1* CRISPR edited strain was to test their behaviour in a motility based assay (Rawsthorne *et al*., 2020). Given that the dumpy phenotype would confound this assessment, we backcrossed the *nlg-1* CRISPR strain with the N2 strain and screened for progeny that retained the *nlg-1* CRISPR mutation and visually lost the dumpy phenotype, appearing phenotypically similar to wild-type in terms of body length and motility (Figure 1D). Sanger sequencing was performed and confirmed the generation of the R451C mutation in *nlg-1* (Figure 1E).

### *unc-58* CRISPR mutation is predicted to result in a non-functional ion channel protein

As well as HDR, CRISPR/Cas9 can also result in NHEJ which can introduce random errors into the DNA sequence in order to repair it (Pannunzio, Watanabe and Lieber, 2018). With this in mind we sequenced the regions surrounding the targeted co-CRISPR loci in order to confirm if the sequences were wild-type as suggested by the lack of *unc-58* or *dpy-10* phenotypes displayed by the CRISPR edited strain (Figure 1D). Thus, we confirmed that the *dpy-10* targeted sequence was wild-type (Figure 2A). However, the targeted *unc-58* sequence was mutated (Figure 2B). We observed a two nucleotide (nt) deletion followed by a 15nt insertion (Figure 2B and C) at the predicted Cas9 cut site (Arribere *et al*., 2014). The disrupted *unc-58* gene causes a frame shift that generates a premature stop codon at position 467, deleting approximately 70% of the predicted C-terminal domain (Figure 2D). Protein sequence analysis also showed that the frameshift mutation alters the amino acid sequence from position 431, which in the WT protein is predicted to be within the fourth transmembrane domain of the potassium channel subunit (Figure 2D) and predicts the mutant UNC-58 protein may have an incomplete fourth transmembrane domain. Overall this suggests that the mutant *unc-58* gene identified in this study is unlikely to encode a functional protein.

**Figure 2:**
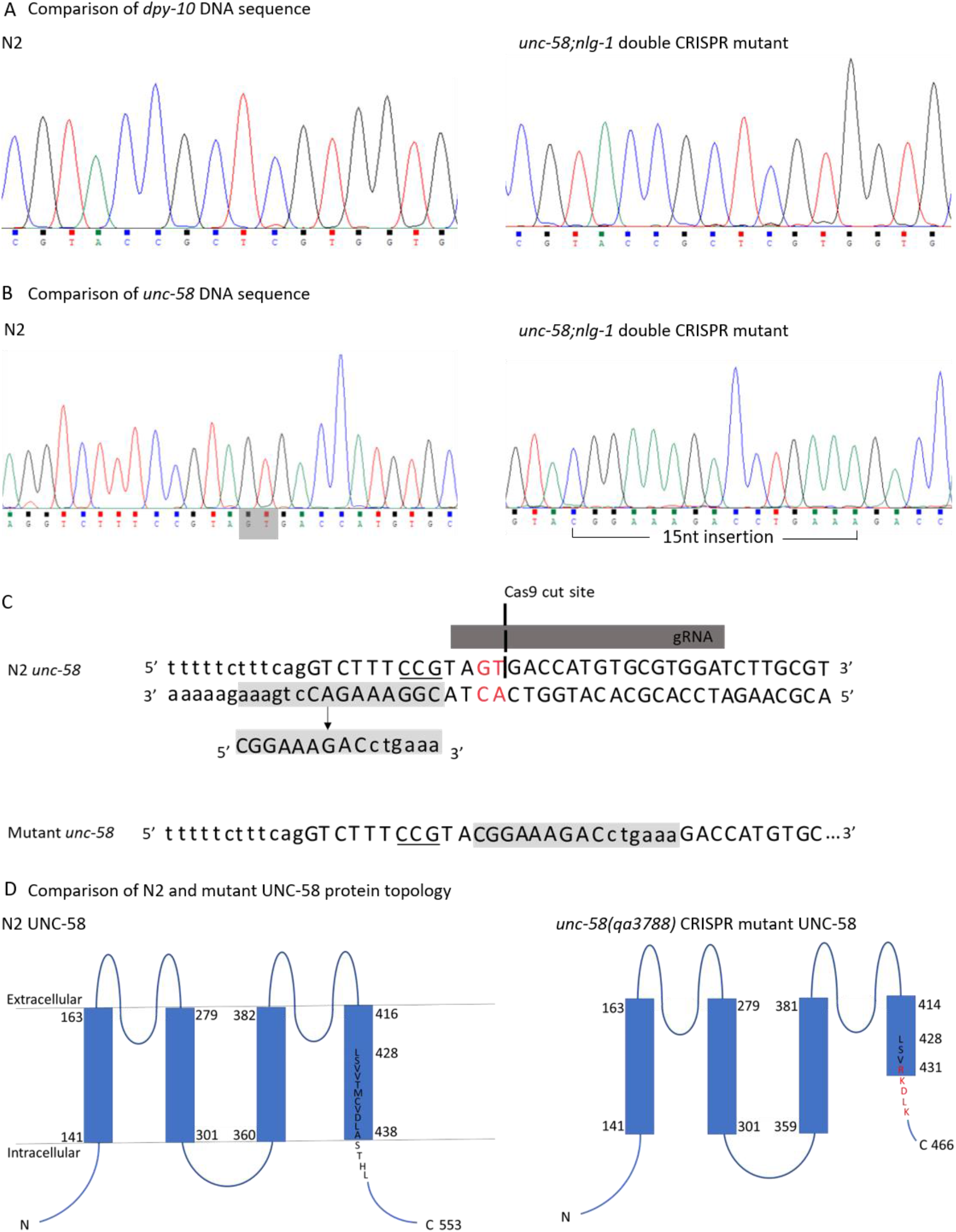
CRISPR mediated mutation to the co-CRISPR gene *unc-58* is predicted to result in a non-functional ion channel protein. **A** Chromatograms comparing partial *dpy-10* DNA sequence from N2 and *unc-58(qa3788);nlg-1(qa3780)* double CRISPR mutant confirming the *dpy-10* sequence is wild-type in both. **B** Chromatograms comparing partial *unc-58* DNA sequence from N2 and *unc-58(qa3788);nlg-1(qa3780)* double CRISPR mutant. Two NTs, GT, that exist in the WT sequence but absent in the mutated *unc-58* sequence are shaded in grey. A 15nt sequence found in the *unc-58(qa3788);nlg-1(qa3780)* double CRISPR mutant is highlighted. All DNA sequences are shown in the 5’ to 3’ orientation. **C** Diagram illustrating the possible origin of the 15nt insert found in the CRISPR mutant. Double stranded wild-type *unc-58* DNA sequence is shown, with the DNA sequence orientation indicated. The gRNA binding site and Cas9 cut site are shown. The PAM sequence is underlined. The two NTs coloured in red indicate the NTs that are deleted as part of the mutation. On the *unc-58* wild-type antisense strand, 15nts are shaded in grey. These NTs correspond to the 15nt insert when read in the 5’ to 3’ orientation. Single stranded mutant *unc-58* sequence containing the 15nt insert is shown. **D** Comparison of wild-type and *unc-58(qa3788)* mutant UNC-58 protein predicted transmembrane domain topology. Wild-type amino acid sequence is shown in black and the mutated amino acid sequence is shown in red.

Discovery of the *unc-58* mutation meant that our CRISPR edited strain, which displayed no obvious *unc-58* marked phenotype, was in fact an *unc-58(qa3788);nlg-1(qa3780)* double mutant strain. Genetic crosses, combined with PCR amplification, enzymatic digest and sequencing allowed the identification of a single strain that retained the *nlg-1* CRISPR mutation and had lost the *unc-58* CRISPR mutation (Figure S1). In the process we also identified strains that retained the *unc-58* CRISPR mutation and lost the *nlg-1* CRISPR mutation.

### Phenotypic analysis of *unc-58* CRISPR mutant identifies it is LOF and effects worm motility

The overarching aim for generating the *nlg-1* CRISPR edited strain was to investigate its effect on the social biology of the worm using a food leaving assay as a paradigm (Rawsthorne *et al*., 2020). Given this, we were interested to understand if the *unc-58* CRISPR mutation, when contemporary to the *nlg-1* CRISPR mutation, would have confounded the assessment of food leaving behaviour. To investigate this, we performed a food leaving assay in which worms were picked onto the centre of a bacterial lawn before food leaving events were quantified at 2 and 24 hours. We have previously shown that N2 worms increase their food leaving rate over 24 hours producing a rate of approximately 0.08 leaving events/worm/minute (Rawsthorne *et al*., 2020). In comparison, both a *nlg-1* null mutant and the *nlg-1* CRISPR mutant showed significantly reduced food leaving rate at 24 hours (Figure 3A) (Rawsthorne *et al*., 2020). Interestingly, the *unc-58;nlg-1* double mutant also showed reduced food leaving behaviour (Figure 3A). In contrast, both the *unc-58* CRISPR mutant and the *unc-58* null mutant showed food leaving behaviour which was similar to the N2 control (Figure 3A). The *unc-58* GOF mutant displays an uncoordinated phenotype (Figure 1B) and we were unable to get a readout of food leaving behaviour (Figure 3A). Taken together, these data show that the *unc-58* CRISPR mutation does not produce a food leaving phenotype and did not confound the assessment of the *nlg-1* CRISPR mutant food leaving behaviour.

**Figure 3:**
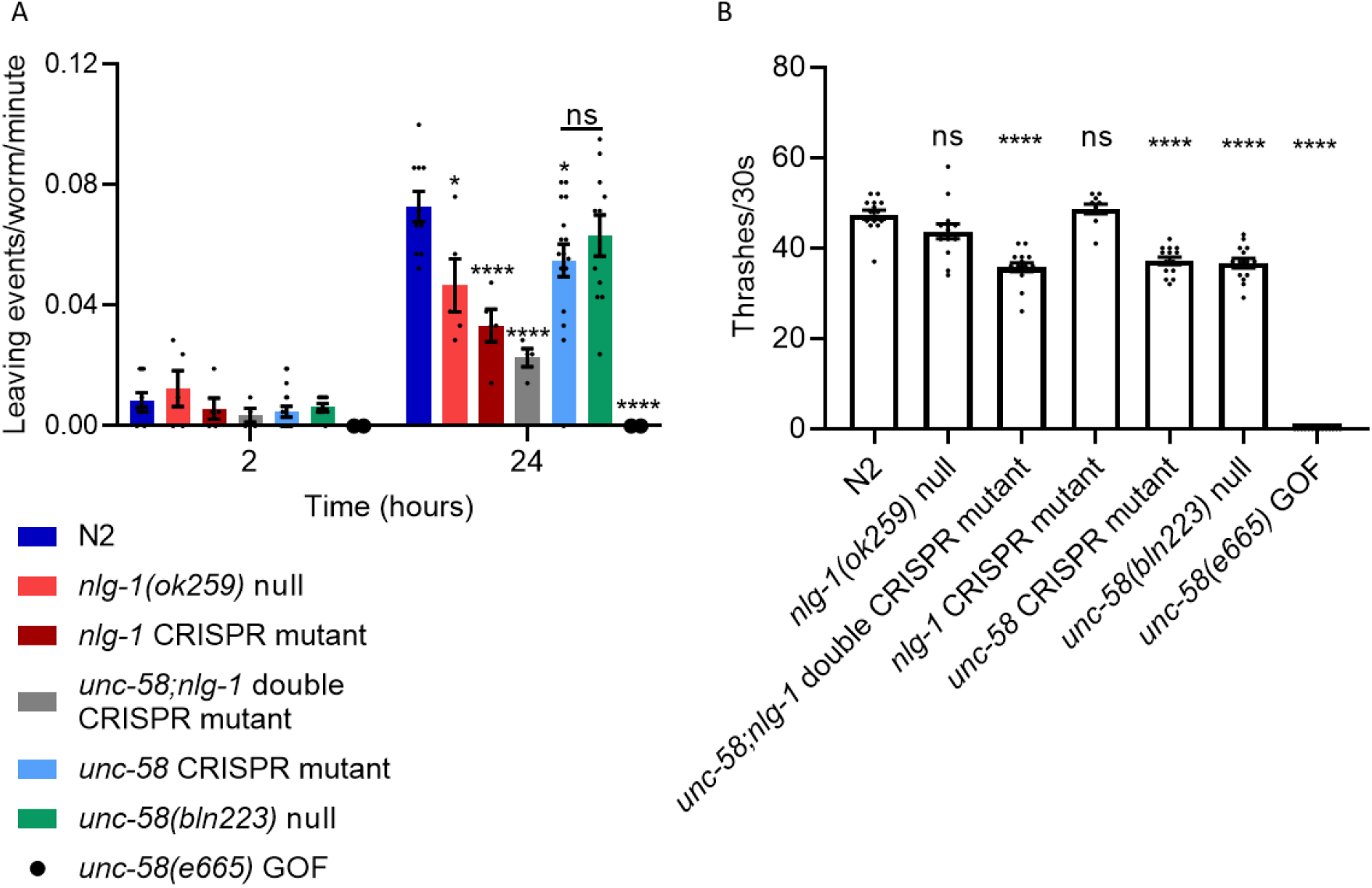
*unc-58* CRISPR mutant phenocopies *unc-58(bln223)* null mutant and is likely to be a LOF mutant. **A** A food leaving assay was performed by picking N2 or mutant worms onto the centre of a bacterial lawn before food leaving events were counted at 2 and 24 hours. *unc-58(qa3788)* CRISPR mutant phenocopies the food leaving behaviour of *unc-58(bln223)* null mutant and N2 at 24 hours. N2 n=10, *nlg-1(ok259)* null n= 5, *nlg-1(qa3780)* CRISPR mutant n=5, *unc-58(qa3788);nlg-1(qa3780)* double mutant (x2 independent lines) n=4, *unc-58(qa3788)* CRISPR mutant (x2 independent lines) n= 16, *unc-58(bln223)* null n= 11, *unc-58(e665)* GOF n=2. Statistical analysis performed using a Two-way ANOVA with Tukey’s multiple comparison test; ns P≥0.05, * P<0.05, **** P≤0.0001. **B** The number of thrashes/30 seconds was counted for N2 and mutant *C. elegans* in liquid medium. The *unc-58(qa3788)* CRISPR mutant shows reduced thrashing behaviour, phenocopying that of *unc-58(bln223)* null mutant. N2 n= 14, *nlg-1(ok259)* n= 14, *unc-58(qa3788);nlg-1(qa378)* double mutant n= 15, *nlg-1(qa3780)* CRISPR mutant n= 10, *unc-58(qa3788)* CRISPR mutant n= 14, *unc-58(bln223)* null n=14, *unc-58(e665)* GOF n=14. Statistical analysis performed using a one-way ANOVA and Dunnett’s multiple comparison test; ns, p>0.05; ****, p≤0.0001. All significance relates to a comparison with N2 control. Data are mean ±SEM.

Given the functional importance of the UNC-58 potassium channel subunit for locomotion (Kasap *et al*., 2018), we wanted to test the intrinsic locomotion of our *unc-58* CRISPR edited strain. To do this we picked worms into liquid medium and observed their thrashing behaviour. We observed that both the *nlg-1* null mutant and the *nlg-1* CRISPR mutant had no thrashing phenotype, thrashing at a similar rate to the N2 control (Figure 3B). In comparison, the *unc-58;nlg-1* double CRISPR mutant, *unc-58* CRISPR mutant and *unc-58* null mutant all showed a similar thrash rate which was significantly lower compared to the N2 paired control (Figure 3B). Furthermore, the *unc-58* GOF mutant did not thrash due to its uncoordinated phenotype (Figure 3B). Taken together, these data show that the *unc-58* CRISPR mutant phenocopies the *unc-58(bln223)* null mutant, suggesting that the CRISPR mediated mutation results in loss of function to the *unc-58* gene. These data also show that the loss of function to *unc-58* in the *unc-58;nlg-1* double mutant results in a reduced thrashing rate which is not displayed in the *nlg-1* CRISPR mutant.

Overall, in the process of CRISPR editing the *nlg-1* gene we unexpectedly created a loss of function mutation within the *unc-58* gene as a consequence of targeting it with reagents designed to generate a gain-of-function phenotype. Analysis of the *unc-58* CRISPR mutant demonstrates that it has a predicted protein sequence that would result in a severe loss of channel function. We show that this loss of function mutation in *unc-58* results in a subtle, but significant effect on motility consistent with its established role in regulation of neuronal membrane potential (Kasap *et al*., 2018).

## Discussion

We demonstrate the use of a previously described method of CRISPR/Cas9 that relies upon the use of co-CRISPR genes (El Mouridi *et al*., 2017) to faciliate screening of transformed worms on the basis of a clearly observable phenotype. The first step in this process involved the transplantation of a d10 sequence in *nlg-1* and utilised the *unc-58* GOF phenotype to facilitate the screening for CRISPR events. The efficiency of d10 transplantation into the gene of interest, *nlg-1*, was lower than that previously reported (El Mouridi *et al*., 2017). The high efficiency of the *dpy-10* sgRNA (El Mouridi *et al*., 2017) meant that fewer worms needed to be injected to produce *dpy-10* marked progeny. However, within the pool of marked progeny the edit efficiency at *nlg-1* was approximately 1%, which is lower than that reported in other genes (El Mouridi *et al*., 2017). This likely reflects that the efficiency varies depending on the gene being targeted.

Our use of *unc-58* as a co-CRISPR gene led to the creation of a novel *unc-58* mutant strain and we have provided evidence that the *unc-58* CRISPR strain phenocopies an *unc-58* null mutant suggesting it is LOF. K2P LOF mutants do not display an easily recognisable phenotype, unlike GOF mutants and are therefore comparitively less well studied (Kasap *et al*., 2018). *unc-58* is expressed widely in interneurons and motor neurons (Salkoff *et al*., 2001). It is hypothesised that GOF to this channel results in loss of ion selectivity and therefore allows the passage of sodium ions. This may then facilitate the depolarisation of motor neurons and may explain the uncoordinated phenotype (Kasap *et al*., 2018) and the hypercontracted state seen in our study and reported more widely (Park and Horvitz, 1986). To our knowledge there is no electrophysiological data to support this hypothesis.

In comparison, it has been suggested that loss-of-function *unc-58* mutants display a less severe phenotype because there is functional redundancy with other K2P channels and this may compensate for the effect of the loss of UNC-58 on resting membrane potential (Kasap *et al*., 2018). Of the few K2P LOF mutants that have been investigated it was shown that the *twk-7* null has moderately enhanced motility (Lüersen, Gottschling and Döring, 2016). This, combined with our findings that the CRISPR mutated *unc-58* is required for normal thrashing behaviour demonstrates that K2P LOF mutants can display locomotory phenotypes, albeit less severe than for GOF mutants. This is consistent with K2Ps playing an important role in co-ordinating normal functioning of the nervous system and highlights how recombination events within this gene could impact downstream analysis of complex integrative behaviours that are dependent upon the worms being able to execute locomotory behaviour.

We show that in the *unc-58* CRISPR LOF mutant the amino acid sequence is mutated in the second half of the fourth transmembrane domain and the majority of the intracellular C-terminus is deleted. This suggests that these domains are likely to play a cruicial role in *unc-58* channel functioning. In mammalian K2Ps the C-terminus is important for channel gating (Piechotta *et al*., 2011). Studies are beginning to define the mechanism of gating in *C. elegans* K2Ps (Ben Soussia *et al*., 2019) however further analysis is needed to ellucidate if there is a conserved role of the C-terminus in channel gating in *C. elegans*.

The caveats of using *unc-58* as a co-CRISPR gene have been previously discussed (Arribere *et al*., 2014). Unlike *unc-58*, other more favourable co-CRISPR genes, such as *dpy-10* and *sqt-1* produce easily recognisable phenotypes in response to both GOF and LOF (Arribere *et al*., 2014). For example, mutation to *dpy-10* can result in dumpy, roller or dumpy roller phenotypes depending on the mutant genotype (Arribere *et al*., 2014). Despite this limitation, *unc-58* continues to be used in co-CRISPR approaches (El Mouridi *et al*., 2017; Wang *et al*., 2018; Schreier *et al*., 2020). We have shown that when using *unc-58* as a co-CRISPR gene, relying soley on the gross phenotypic appearance of the worm is not a robust way to assume genotype. Therefore, we suggest that sequencing of co-CRISPR genes, particularly *unc-58*, is important following their use.

In conclusion, the visibly obvious hypercontraction in the viable adult *C. elegans* that is generated by a GOF mutation in *unc-58* provides an excellent marker for screening for CRISPR events in target genes of interest. In using this approach we unexpectidely generated a novel *unc-58* mutation and provided evidence that this LOF impaired motility in liquid. This will facilitate the understanding of K2Ps in *C. elegans* and the behaviours they regulate. However, it is clear that if utilised to generate mutants for investigation of more complex behaviours or readouts that depend on neuronal exciatbility, the spurious re-arrangements at the *unc-58* marker locus can generate more subtle but not inconsequential LOF mutations. Based on our experience of using *unc-58* in co-CRISPR strategies destined to target gene loci that impact discrete determinants of integrative behaviour, we conclude that this requires routine sequencing of the co-CRISPR gene to avoid potential confounds of unpredicted CRISPR events.

## References

Adli, M. (2018) ‘The CRISPR tool kit for genome editing and beyond’, Nat Commun, 9(1), pp. 1911.

Arribere, J. A., Bell, R. T., Fu, B. X. H., Artiles, K. L., Hartman, P. S. and Fire, A. Z. (2014) ‘Efficient Marker-Free Recovery of Custom Genetic Modifications with CRISPR/Cas9 in Caenorhabditis elegans’, Genetics, 198(3), pp. 837–U842.

Ben Soussia, I., El Mouridi, S., Kang, D., Leclercq-Blondel, A., Khoubza, L., Tardy, P., Zariohi, N., Gendrel, M., Lesage, F., Kim, E.-J., Bichet, D., Andrini, O. and Boulin, T. (2019) ‘Mutation of a single residue promotes gating of vertebrate and invertebrate two-pore domain potassium channels’, Nature Communications, 10(1), pp. 787.

Brenner, S. (1974) ‘The genetics of Caenorhabditis elegans’, Genetics, 77(1), pp. 71–94.

Dickinson, D. J. and Goldstein, B. (2016) ‘CRISPR-Based Methods for Caenorhabditis elegans Genome Engineering’, Genetics, 202(3), pp. 885–901.

El Mouridi, S., Lecroisey, C., Tardy, P., Mercier, M., Leclercq-Blondel, A., Zariohi, N. and Boulin, T. (2017) ‘Reliable CRISPR/Cas9 Genome Engineering in Caenorhabditis elegans Using a Single Efficient sgRNA and an Easily Recognizable Phenotype’, G3-Genes Genomes Genetics, 7(5), pp. 1429–1437.

Greene, J. S., Dobosiewicz, M., Butcher, R. A., McGrath, P. T. and Bargmann, C. I. (2016) ‘Regulatory changes in two chemoreceptor genes contribute to a Caenorhabditis elegans QTL for foraging behavior’, Elife, 5.

Hartman, P. S., Barry, J., Finstad, W., Khan, N., Tanaka, M., Yasuda, K. and Ishii, N. (2014) ‘Ethyl methanesulfonate induces mutations in Caenorhabditis elegans embryos at a high frequency’, Mutat Res, 766-767, pp. 44-8.

He, F. (2011) ‘Single Worm PCR’, Bio-protocol, 1(8), pp. e60.

Hsu, P. D., Lander, E. S. and Zhang, F. (2014) ‘Development and applications of CRISPR-Cas9 for genome engineering’, Cell, 157(6), pp. 1262–1278.

Huumonen, K., Immonen, H. K., Baverstock, K., Hiltunen, M., Korkalainen, M., Lahtinen, T., Parviainen, J., Viluksela, M., Wong, G., Naarala, J. and Juutilainen, J. (2012) ‘Radiation- induced genomic instability in Caenorhabditis elegans’, Mutat Res, 748(1-2), pp. 36–41.

Jamain, S., Quach, H., Betancur, C., Rastam, M., Colineaux, C., Gillberg, I. C., Soderstrom, H., Giros, B., Leboyer, M., Gillberg, C., Bourgeron, T. and Paris Autism Res Int Sibpair, S. (2003) ‘Mutations of the X-linked genes encoding neuroligins NLGN3 and NLGN4 are associated with autism’, Nature Genetics, 34(1), pp. 27–29.

Jinek, M., Chylinski, K., Fonfara, I., Hauer, M., Doudna, J. A. and Charpentier, E. (2012) ‘A Programmable Dual-RNA-Guided DNA Endonuclease in Adaptive Bacterial Immunity’, Science, 337(6096), pp. 816–821.

Kaletta, T. and Hengartner, M. O. (2006) ‘Finding function in novel targets: C-elegans as a model organism’, Nature Reviews Drug Discovery, 5(5), pp. 387–398.

Kasap, M. and Dwyer, D. S. (2020) ‘Clozapine, nimodipine and endosulfan differentially suppress behavioral defects caused by gain-of-function mutations in a two-pore domain K(+) channel (UNC-58)’, Neurosci Res.

Kasap, M., Weeks, K., Aamodt, E. J. and Dwyer, D. S. (2018) ‘Gain-of-function mutations in a two-pore domain K+ channel (unc-58) cause developmental, motor and feeding defects in C. elegans modified by temperature and a channel inhibitor, loratadine’, Current Neurobiology, 9(3), pp. 84–91.

Kim, H., Ishidate, T., Ghanta, K. S., Seth, M., Conte, D., Shirayama, M. and Mello, C. C. (2014) ‘A Co-CRISPR Strategy for Efficient Genome Editing in Caenorhabditis elegans’, Genetics, 197(4), pp. 1069–U37.

Krogh, A., Larsson, B., von Heijne, G. and Sonnhammer, E. L. (2001) ‘Predicting transmembrane protein topology with a hidden Markov model: application to complete genomes’, J Mol Biol, 305(3), pp. 567–80.

López-Cruz, A., Sordillo, A., Pokala, N., Liu, Q., McGrath, P. T. and Bargmann, C. I. (2019) ‘Parallel Multimodal Circuits Control an Innate Foraging Behavior’, Neuron, 102(2), pp. 407- 419.e8.

Lüersen, K., Gottschling, D.-C. and Döring, F. (2016) ‘Complex Locomotion Behavior Changes Are Induced in Caenorhabditis elegans by the Lack of the Regulatory Leak K+ Channel TWK- 7’, Genetics, 204(2), pp. 683–701.

Ma, D. and Liu, F. (2015) ‘Genome Editing and Its Applications in Model Organisms’, Genomics, proteomics & bioinformatics, 13(6), pp. 336–344.

McDiarmid, T. A., Belmadani, M., Liang, J., Meili, F., Mathews, E. A., Mullen, G. P., Hendi, A., Wong, W.-R., Rand, J. B., Mizumoto, K., Haas, K., Pavlidis, P. and Rankin, C. H. (2019) ‘Systematic phenomics analysis of autism-associated genes reveals parallel networks underlying reversible impairments in habituation’, Proceedings of the National Academy of Sciences, pp. 201912049.

Mojica, F. J., Díez-Villaseñor, C., García-Martínez, J. and Soria, E. (2005) ‘Intervening sequences of regularly spaced prokaryotic repeats derive from foreign genetic elements’, J Mol Evol, 60(2), pp. 174–82.

Paix, A., Folkmann, A., Rasoloson, D. and Seydoux, G. (2015) ‘High Efficiency, Homology- Directed Genome Editing in Caenorhabditis elegans Using CRISPR-Cas9 Ribonucleoprotein Complexes’, Genetics, 201(1), pp. 47-+.

Paix, A., Wang, Y. M., Smith, H. E., Lee, C. Y. S., Calidas, D., Lu, T., Smith, J., Schmidt, H., Krause, M. W. and Seydoux, G. (2014) ‘Scalable and Versatile Genome Editing Using Linear DNAs with Microhomology to Cas9 Sites in Caenorhabditis elegans’, Genetics, 198(4), pp. 1347-+.

Pannunzio, N. R., Watanabe, G. and Lieber, M. R. (2018) ‘Nonhomologous DNA end-joining for repair of DNA double-strand breaks’, The Journal of biological chemistry, 293(27), pp. 10512–10523.

Park, E. C. and Horvitz, H. R. (1986) ‘Mutations with dominant effects on the behavior and morphology of the nematode Caenorhabditis elegans’, Genetics, 113(4), pp. 821–852.

Piechotta, P. L., Rapedius, M., Stansfeld, P. J., Bollepalli, M. K., Ehrlich, G., Andres-Enguix, I., Fritzenschaft, H., Decher, N., Sansom, M. S. P., Tucker, S. J. and Baukrowitz, T. (2011) ‘The pore structure and gating mechanism of K2P channels’, The EMBO journal, 30(17), pp. 3607–3619.

Rawsthorne, H., Calahorro, F., Feist, E., Holden-Dye, L., O’Connor, V. and Dillon, J. (2020) ‘Neuroligin dependence of social behaviour in C. elegans provides a model to investigate an autism associated gene’, Human Molecular Genetics.

Reiner, D. J., Weinshenker, D. and Thomas, J. H. (1995) ‘Analysis of dominant mutations affecting muscle excitation in Caenorhabditis elegans’, Genetics, 141(3), pp. 961–976.

Salkoff, L. (2005) Potassium channels in C. elegans The C. elegans Research Community, WormBook.

Salkoff, L., Butler, A., Fawcett, G., Kunkel, M., McArdle, C., Paz-y-Mino, G., Nonet, M., Walton, N., Wang, Z. W., Yuan, A. and Wei, A. (2001) ‘Evolution tunes the excitability of individual neurons’, Neuroscience, 103(4), pp. 853–9.

Schreier, J., Dietz, S., de Jesus Domingues, A. M., Seistrup, A.-S., Nguyen, D. A. H., Gleason, E. J., Ling, H., L’Hernault, S. W., Phillips, C. M., Butter, F. and Ketting, R. F. (2020) ‘A membraneassociated condensate drives paternal epigenetic inheritance in <em>C. elegans</em>’, bioRxiv, pp. 2020.12.10.417311.

Sonnhammer, E. L., von Heijne, G. and Krogh, A. (1998) ‘A hidden Markov model for predicting transmembrane helices in protein sequences’, Proc Int Conf Intell Syst Mol Biol, 6, pp. 175–82.

Sonnhammer, E. L. L. and Durbin, R. (1997) ‘Analysis of protein domain families in Caenorhabditis elegans’, Genomics, 46(2), pp. 200–216.

Wang, H., Park, H., Liu, J. and Sternberg, P. W. (2018) ‘An Efficient Genome Editing Strategy To Generate Putative Null Mutants in Caenorhabditis elegans Using CRISPR/Cas9’, G3 (Bethesda, Md.), 8(11), pp. 3607–3616.

Wong, W. R., Brugman, K. I., Maher, S., Oh, J. Y., Howe, K., Kato, M. and Sternberg, P. W. (2019) ‘Autism-associated missense genetic variants impact locomotion and neurodevelopment in Caenorhabditis elegans’, Human Molecular Genetics, 28(13), pp. 2271–2281.

