## Supplementary figure for "Confounds of using the *unc-58* selection marker highlights the importance of genotyping co-CRISPR genes"

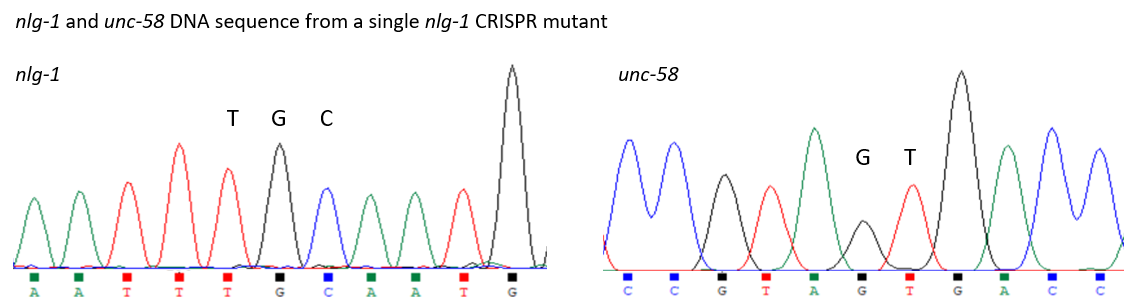

Figure S1: Chromatograms showing partial *nlg-1* and *unc-58* DNA sequence confirming the *nlg-1(qa3780)* CRISPR mutant carries the R451C mutation as well as wild-type *unc-58* sequence. The TGC codon that encodes the R451C mutation is indicated. The wild-type GT NTs that were deleted as part of the *unc-58* CRISPR mutation are indicated. DNA sequences are shown in the 5’ to 3’ orientation.
